# Protein synthesis is suppressed in sporadic and familial Parkinson’s Disease by LRRK2

**DOI:** 10.1101/2020.04.27.053694

**Authors:** Prasannakumar Deshpande, Dani Flinkman, Ye Hong, Elena Goltseva, Valentina Siino, Lihua Sun, Sirkku Peltonen, Laura Elo, Valtteri Kaasinen, Peter James, Eleanor T. Coffey

## Abstract

Gain of function LRRK2-G2019S is the most common mutation associated with both familial and sporadic Parkinson’s disease. It is expected therefore that understanding the cellular function of LRRK2 will provide much needed insight on the pathological mechanism of sporadic Parkinson’s, which is the most common form. Here we show that constitutive LRRK2 activity represses nascent protein synthesis in rodent neurons. Specifically, enzymatic inhibition of LRRK2, gene silencing or gene knockout of *Lrrk2* increase translation. In the rotenone model of Parkinson’s, LRRK2 activity increases, leading to repression of translation and dopaminergic neuron atrophy both of which are prevented by LRRK2 inhibition. This is accompanied by altered phosphorylation of eIF2α-S52(↑), eIF2s2-S2(↓) and eEF2-T57(↑) in striatum/substantia nigra in a direction that signifies inhibition of protein synthesis. Significantly, LRRK2 is activated and translation is 40% reduced in Parkinson’s patient fibroblasts (G2019S and sporadic) and LRRK2 inhibition restores normal translation. In contrast, translation is unchanged in cells from multiple system atrophy patients, implying disease specificity. These findings suggest that LRRK2-dependent repression of translation may be a proximal function of LRRK2 in Parkinson’s pathology.

## INTRODUCTION

Parkinson’s disease is the second most common neurodegenerative disorder after Alzheimer’s disease with the highest prevalence in Europe and North and South America (1, 2). No cure exists and long term symptomatic treatment is associated with complications. Although its neuropathological basis is not fully understood, degeneration of dopaminergic neurons in the *substantia nigra pars compacta* is repeatedly shown, and subsequent reduction of dopamine release in the basal ganglia leads to Parkinsonian motor symptoms (2). Meta-analysis of genes associated with Parkinson’s disease has revealed over 40 risk loci (3). Among these, leucine rich-repeat kinase 2 (*LRRK2*)-*G2019S* is the most common mutation and it is associated with late onset Parkinson’s disease. *LRRK2*-G2019S confers a gain of function in kinase activity (4), as do disease-associated mutations in the GTPase domain(5–7). In addition to its association with familial Parkinson’s, gene wide association studies have highlighted *LRRK2* as a risk factor for sporadic Parkinson’s (8). Because of this and the finding that *LRRK2-G2019S* cases are symptomatically indistinguishable from sporadic ones (9–11), LRRK2 may also contribute to the pathology of sporadic Parkinson’s (12). Consequently, identifying the cellular function of LRRK2 is expected to help elucidate the pathological mechanism of sporadic Parkinson’s disease, which accounts for around 90% of all cases.

A large body of evidence suggests that LRRK2 regulates endolysosomal trafficking and that this may contribute to the pathology of Parkinson’s disease (13–15). In addition to this however, LRRK2 has also been implicated in the regulation of protein synthesis through the phosphorylation of ribosomal proteins and regulators of ribosomal function (16–19). These studies suggest that LRRK2 augments RNA translation. However this is in contrast to clinical data where increased phosphorylation of eIF2α and eEF2 on sites that signal protein synthesis arrest was detected in substantia nigra of Parkinson’s patients (20, 21). Also, expression of implicated regulators of endolysosomal trafficking (VPS35, Gcase1, LAMP2A and ATP13A2) are decreased in post-mortem brains from Parkinson’s patients carrying LRRK2 mutations (13, 15). Despite the supporting clinical data, the possibility that LRRK2 could suppress translation has not received much attention.

Here we took a basic discovery approach to identify what type of brain organelle was most highly phosphorylated by the disease-associated LRRK2-G2019S. This highlighted ribosomal fractions as preferentially phosphorylated and this led us examine whether endogenous LRRK2 regulated RNA translation. We found that translation was decreased in both the rotenone and 6-hydroxy dopamine models of Parkinson’s disease and in patient fibroblasts. Moreover, when we examined cells from sporadic and G2019S patients, we also found that translation was decreased and reversed upon LRRK2 inhibition. A shot-gun phospho-proteome analysis combined with antibody approach revealed that checkpoint regulators of protein synthesis arrest were switched on in Parkinson’s models and in cells from sporadic patients. This data implies that protein synthesis is repressed in sporadic Parkinson’s disease by elevated LRRK2.

## MATERIALS *AND* METHODS

### Antibodies, tools

Antibodies for LRRK2 [MJFF2 (C41-2)] (# ab133474), phospho-S935-LRRK2 [UDD2 10 (12)] (#ab133450), phospho-T73-Rab10 [MJF-R21] (# ab230261), Rab10 [MJF-R23] (# ab237703), and RPL10a (# ab55544) were from Abcam. Anti-tyrosine hydroxylase was from Millipore (#AB1542). Antibodies for eIF2B5(sc-28854), eIF2α (sc-11386), phospho-eIF2α–S52 (sc-101670), eIF4G3 (sc-100732) and Ndufs3 (sc-292169) were from Santa Cruz Biotechnology. Anti-Rab8 (#R66320) and Rab4 (#R68520) were from BD Transduction Laboratories. Anti-4E-BP1 (# 9452), eEF2 (#2332) and phospho-eEF2(T57) (#2331) were from Cell Signaling Technology and anti-actin (# A3853) was from Sigma-Aldrich. LRRK2-IN1 was from Merck Millipore (# 438193), GSK-2578215A (#4629) and MLi-2 (# 5756) were from Tocris Bioscience. Miglyol 812N was from Cemer Oleo, GmbH and Co, KG (Germany). Patient cells were from the National Institute of Neurological Disorders and Stroke (NINDS) repository at the Coriell Institute for Medical Research and the Cell Line and DNA Biobank from Patients Affected by Genetic Diseases, Telethon Network of Genetic Biobanks (TNGB) (project no. GTB12001) (22). C57BL/6-*Lrrk2^tm1.1Mjff^*/J mice developed by Michael J. Fox Foundation (23) were obtained from The Jackson Laboratory (Stock no. 016121).

### Plasmid

The bicistronic CAP/IRES reporter(pYIC) was a gift from Han Htun (24).

### Separation of mitochondrial and ribosomal fractions

Mitochondrial and ribosomal fractions were isolated from rat brain as previously (25). Briefly, rat brains were homogenized in 12 ml homogenization buffer (300 mM sucrose, 10 mM HEPES, pH7.4, 0.1 mM EGTA, 2 mM EDTA) with protease inhibitors (1 μg/ml leupeptin, pepstatin and aprotinin and 100 μg/ml PMSF). Lysate was kept on ice for 20 min and unbroken cells (P1) were removed by centrifugation at 2800 × g_avg_ for 10 min at 4°C. Supernatant 1 (S1) was centrifuged at 22,000 × g_avg_ at 4°C for 10 min using a SW41Ti rotor and Beckman L90K ultracentrifuge. The subsequent pellet (P2) contained mitochondria. The remaining supernatant 2 (S2) was centrifuged at 100,000 × gavg at 4°C for 8 h to give pellet 3 (P3) which contained ribosomes.

### Sucrose gradient fractionation of ribosomal subunits

P14 rat brain was homogenized with a Potter-Elvehjem homogenizer (20 strokes) in 5 ml of ice-cold homogenization buffer (5 % sucrose, 10 mM HEPES, pH7.4, 0.1 mM EGTA, 2 mM EDTA) with protease inhibitors (as above) and the second supernatant (S2) fraction was prepared as before. S2 was loaded on a linear sucrose gradient (5-50% in 8 ml). The gradient was centrifuged at 150,000 × g at 4°C in a SW41Ti rotor for 2 h and 40 min. Fractions of 0.5 ml were collected from the bottom of the tube using a peristaltic pump and RNA amount was determined by measuring absorbance at 260 nm.

### In vitro phosphorylation of brain fractions using LRRK2-G2019S

The assay was carried out as previously with modifications (26). Briefly, the P3 ribosomal pellet or the P2 mitochondrial pellet from P7 rat brain were resuspended in equal volume of buffer (20 mM MOPS, pH 7.2, 2 mM EGTA, 2 mM EDTA, 10 % glycerol, 1 mM benzamidine) with protease inhibitors (1 μg/ml leupeptin, pepstatin and aprotinin and 100 μg/ml PMSF) and 1 % IGEPAL. After 5 min on ice, the lysate was homogenized with 8 strokes using a 27G syringe and then centrifuged at 16,000 × g at 4°C. 0.1 mM ZnCl2 and 2.5 U Antarctic phosphatase (New England Biolabs) were added to the supernatant and incubated at 37°C for 2 h followed by heat inactivation of phosphatase at 65°C, 10 min. Lysates were desalted using Micro Biospin columns from Biorad (catalogue # 732-6223). To start the kinase reaction, 5 μCi of γ-[_32_P] ATP, MgCl_2_ (10 mM) and 44.12 nM LRRK2-G2019S (Invitrogen, cat# PV4881) was added. Reactions were carried out at 30°C for 1 h and ended by adding *Laemmli* buffer. Radioactive samples were separated by SDS-PAGE and visualized by autoradiography.

### Hippocampal and midbrain cultures

Hippocampal neurons were isolated from new born Sprague Dawley rats as previously described (27). *Lrrk2* knockout and wild-type neurons were litter matched from *Lrrk2+/−* breeding pairs. Genotyping was done on cell lysates following Jackson Laboratory recommendations. Midbrain cultures were isolated from P0 Sprague Dawley rats or C57BL6 mice. Midbrains in dissection medium (30 mM K_2_SO_4_, 81.8 mM Na_2_SO_4_, 5.8 mM MgCl_2_, 1 mM D-glucose, 0.25 mM Hepes pH 7.4, 0.001% Phenol red, supplemented with 1μM kynurenic acid) were digested with Papain (10U/ml Worthington, 3119) as previously described (27). Plating density is indicated in subsequent paragraphs. Cells were maintained in Neurobasal A (Gibco™, Life Technologies) supplemented with B27, 2mM Gln and penicillin 50U/ml and streptomycin 50 μg/ml.

### Patient skin cell isolation and maintenance

Skin punches from patients and healthy donors were isolated at Turku University Hospital (TUH) from the upper arm of individuals by a licensed dermatologist, and placed immediately in *Minimal essential medium* (M.E.M., Sigma Aldrich). Tissue was chopped into several small pieces using a sterile blade and incubated in M.E.M. supplemented with Gln (2 mM), penicillin (50 U/ml) and streptomycin (50 μg/ml) at 37°C, 5% CO_2_. Fibroblasts were passaged when 95% confluent. Experiments were done at passage 7 to 9.

### Neurite atrophy

Midbrain cultures (200,000 cells/well in 96 well plate) were treated at 3 d post-plating with rotenone in DMSO +/− LRRK2 inhibitors (IN1, GSK2578215A or MLi-2) as indicated in the figures. After 24 h, cells were fixed with 4 % PFA in PBS. To identify dopaminergic neurons, tyrosine hydroxylase (TH) was detected by immunostaining as previously described (28). Cells were imaged with a Zeiss TIRF-3 microscope equipped with CMOS Orca Flash 4 (Hamamatsu). All TH-positive cells were counted from tiled images of wells. The number of TH-positive cells that had intact neurites (> 2x soma diameter) were counted and expressed as a % of all TH-positive cells.

### Pyknosis

Neuronal pyknosis was evaluated from TH-positive cells using 4 μg/ml Hoechst-33342. Tile images of nuclei and TH fluorescence were acquired using a Zeiss AxioVert 200M microscope and 10x objective. All TH-positive neurons were scored by a blinded experimenter. Cells with shrunken, bright nuclei were considered pyknotic.

### 6-OHDA treatment

Freshly prepared 6-OHDA (40 mM) in DMSO was added to 3 d *in vitro* midbrain cells plated at 150,000 cells/well in a 96 well plate. First, 180 μl of fresh medium was added. Then 6-OHDA was added to a final concentration of 40 μM with 0.2 % ascorbic acid in 20μl medium. After 15 min, cells were washed 1x with medium and 50 nM MLi-2 or 0.1% DMSO was added. After 24 h, AHA labeling was performed and cells were stained for TH to enable analysis of protein synthesis in dopaminergic neurons.

### Measurement of translation using AHA

was carried out as previously, with modifications (29). Briefly, hippocampal, midbrain or skin cells were plated at densities of 100,000, 500,000 or 50,000 cells per well on round coverslips (1.3 cm diameter). At 20 d *in vitro* (hippocampal, 3 d (dopaminergic) or 48 hours post-plating (skin), cells were washed 1x and incubated with Met-free DMEM (Gibco, Thermo Fisher Scientific) supplemented with 2 mM Gln for 30 min including LRRK2 inhibitors or 0.1% DMSO, followed by Met-free media containing 1 mM L-azidohomoalanine (AHA) + LRRK2 inhibitors/DMSO. Labeling was stopped by washing with 1 ml PBS followed by fixation in 4 % paraformaldehyde. Cells were permeablized overnight in *block* (0.25 % Triton X100, 0.2 % BSA, 5 % sucrose, 10 % horse serum in PBS), and washed 5x with 1 ml of 3 % BSA in PBS. Cycloaddition of Alexa-488 was carried out using the Thermo Fisher Scientific kit (catalogue # A10267) according to the manufacturer’s instructions. Nuclei were stained with 1:2000 Hoechst-33342 in PBS. Coverslips were divided into 4 quadrants and 3-5 fluorescent images were acquired per quadrant using a 40x objective and a Leica DMRE microscope with an ORCA C4742-95 CCD camera (Hamamatsu). Mean fluorescence intensity r.o.i.s in the soma were measured from all cells using ImageJ. Background was determined from identical r.o.i. from cells that did not receive AHA. For dendrites, line intensity measurements were taken from 10 μm lines in primary dendrites. Image acquisition time and excitation power was constant throughout and performed blind. Specifically, slide labels were masked by an uninvolved researcher and assigned code numbers which were only revealed upon completion of analysis.

### Measurement of translation using ^[35]^S-Met

Hippocampal neurons (50,000 cells/well on 48 well plates) were depleted of Met in Met-free DMEM supplemented with 2 mM L-Gln at 20 d post plating for 30 min +/− anisomycin or MLi-2 as indicated. ^[35]^S-Met was added to a final concentration of 200 μCi/ml, for 60 min. After labeling, cells were lysed in *Laemmli* buffer and proteins separated by SDS-PAGE. ^[35]^S-Met incorporation was detected using a Fuji phosphorimager and intensities quantified using ImageJ for the entire gel lane of proteins and normalized to coomassie blue.

### Measurement of translation in a reconstituted assay

A human, cell-free protein expression system from Takara Bio Inc. (Japan) was used according to the manufacturer’s instructions. Briefly a β-galactosidase reporter (150 ng) was incubated with 60 nM GST-G2019S-LRRK2 or GST alone for 1.5 h at 32°C. β-galactosidase expression was measured using colorimetric detection (Abs_420nm_, using a Synergy H1 plate reader (Biotek) from samples in a 96-well plate. Absorbance at 420 nm was measured using the Synergy H1 Hybrid Reader (Biotek).

### LRRK2 knockdown

6 d *in vitro* hippocampal neurons on coverslips were transfected with 200 nM non-targeting (5’-GCUAAUACCUAUCAAUUGUU-3’) or *Lrrk2* siRNA (5’-AAGUUGAUAGUCAGGCUGAAU-3’) (30) and protein synthesis was measured using AHA labelling at 20 d *in vitro*. The efficiency of knockdown was assessed by immunostaining using the LRRK2 antibody [MJFF2 (C41-2)] (# ab133474, Abcam) at1:200 dilution, and secondary detection was with goat anti-Rabbit IgG (H+L)-Alexa Fluor 568 (#A11011, ThermoFisher Scientific) at 1:500 dilution. Fluorescence intensity of regions of interest in soma and dendrites was measured as described above by a blinded experimenter.

### Rotenone treatment of rats and behavioral testing

Group housed female, 12 week old Sprague-Dawley rats were injected intraperitoneally with rotenone (n = 6) (1 mg/kg) diluted in carrier (98 % Miglyol 812N, 2 % DMSO), or with carrier alone (n = 6). Postural instability was tested at 6 and 11 d according to a previous report (31). Briefly, while holding the animal vertically, one forelimb was immobilized to the chest while the other forelimb touched the surface of scored sandpaper. The center of gravity of the animal was advanced until a step was triggered. The average distance taken to trigger a step was measured from both forelimbs. The test was performed between 9 and 11 am before the start of injections (0 d), mid-way (6 d) and one day before termination (11 d) by a blinded experimenter. On 12d, animals underwent terminal anesthesia (150 mg/kg of Mebunat intraperitoneal; Orion Corporation). Brains were rapidly isolated and cut into 1 mm thick coronal sections starting rostral to the optic chiasm and continuting until midway through the pons, using a pre-chilled rat brain matrix (Zivic Instruments) as described previously (32). Sections were frozen on dry ice and striatum and substantia nigra were manually dissected using a scalpel (33). Tissues were denatured using Denator (Denator, Uppsala, Sweden) before storage at −80 °C.

### Phosphopeptide enrichment and mass spectrometry sample analysis

For analysis of phosphoprotein changes in substantia nigra and striatum of rotenone treated rats, animals were treated exactly as described above. Chemicals for digestion and mass spectrometry analysis were from Sigma-Aldrich (St. Louis, USA) unless otherwise stated. Substantia nigra or striatum from rotenone injected (n = 14) or control (n = 12) rats were run 1 cm into a SDS-PAGE, and stained using GelCode blue (Thermo Fisher Scientific). Concentrated samples were excised from the gel, cut in 1 mm^3^ pieces, and destained with 40 mM Ambic/50% for 3 × 15 min. Samples were reduced with 20 mM DTT for 30 min at 56°C, and alkylated with 55 mM Iodoacetoamide for 20 min at RT in the dark. Trypsin digestion was performed overnight with sequencing grade modified Trypsin (Promega, Madison, Wisconsin, USA). Peptides were extracted with 100% ACN and 50% ACN/ 5 % HCOOH, and dried using a SpeedVac for ~2 h. Dried peptides were resuspended in 10 μl 0.1% TFA, followed by addition of 40 μl of 6 % TFA 80 % ACN and loaded on a column consisting of 1 mg of 20 μM, 300Å TiO2 particles (ZirChrom, Anoka, MN, USA) by centrifugation at 3000 rpm for 45 s. Columns were washed twice with 50 μl of 6% TFA/ 80% ACN, and twice with 50 μl of 0.1% TFA. Centrifugation was at 3000 rpm for 30 s between washes. Peptides were eluted with 50 μl of 5% NH_4_OH, and acidified with 100 μl of 10% HCOOH andcleaned using in-house prepared C18 spin columns, and dried with a SpeedVac at RT for ~30 min. Enriched phosphopeptides were resuspended in 15 μl of 0.1% HCOOH, and analysis was carried out using an LC-ESI-MS/MS with an Easy-nLC1200 coupled to an Orbitrap Fusion™ Lumos™ (Thermo Fisher Scientific) operated in Data Dependent Acquisition mode with 3 s cycle time and EThcD collision. Survey scans were acquired at 120K resolution at 200 m/z with 375-1500 m/z scan range, and MS/MS at 30K resolution at 200 m/z. Peptides were first loaded on a 100 μM × 2 cm trapping column, and separated on 75 μM ID × 15 cm analytical column, which were packed with ReproSil-Pur 5 μm 200 Å C18-AQ (Dr. Maisch HPLC GmbH, Ammerbuch-Entringen, Germany). Mobile phase A consisted of 0.1% FA, and B 80/20 (v/v) ACN/water, and peptides were eluted over a 60 min gradient at 300 nl/min.

### Data analysis from mass spectrometry output

MS/MS data analysis was carried out using MaxQuant v1.6.3.4 (34) with *“match between runs”* and *“known contaminants”* selected. Data was searched against *Rattus norvegicus* UniprotKB database. Trypsin/P with 2 allowed miss-cleavages was used and FDR was set at 1% for proteins and peptides. Decoy-target FDR estimation in MaxQuant was used to determine false positives. Allowed modifications were fixed carbamidomethylation at C, variable M oxidation, STY phosphorylation and protein N-terminal acetylation. Data was log2 transformed and the resulting data was analyzed using Perseus v1.6.2.3 (35). Perseus-marked potential contaminants and phosphosites with more than 50% missing values in both rotenone and control conditions were removed. Samples were normalized by subtracting the sample median intensity from individual phosphosite intensities as previously described (35, 36), and the global median of all samples was added to each phosphosite intensity to bring phosphosite intensities back to their original scale. The Perseus *“replace missing values from normal distribution”* function was used. MS/MS statistical analysis used Student’s t-test (p < 0.05) with the Perseus *“permutation based FDR for multiple hypothesis testing correction”* set at FDR < 0.05. A threshold for significantly changing phosphosites of Log2 fold change of 1 was also imposed.

### Western blotting

After treatment, mouse hippocampal (DIV20) or midbrain (DIV8) neurons were lysed in 1x Laemmli (Biorad #1610747) supplemented with 1x protease inhibitor cocktail (#005892791001, Roche) and 2x phosphatase inhibitor cocktail (# P2850, Sigma), 2 mM MgCl_2_ and 25U/ml Benzonase (# 70746, Merck Millipore). Lysates were resolved on 4–20% Mini-PROTEAN® TGX™ Precast Protein Gels (#4561096, Biorad) Membranes or gels were cut in horizontal strips according to defined molecular weight and blocked with 5% milk in Tris buffered saline with 0.01% tween (TBST) for 1 h and probed with primary antibodies overnight at 4°C. Antibodies were used at the following dilutions, phospho-S935 LRRK2 (1:1000), LRRK2 (1:1000), RPL10a (1:10000), 4E-BP1 (1:1000), phospho-4EBP (1:1000), eIF2B5 (1:500), NDUFS3 (1:500), Rab10 (1:3000), phospho-T73-Rab10 (1:1000), eIF2α (1:500), phospho-S52-eIF2α (1:500), phospho-T57-eEF2 (1:1000), eEF2 (1:1000) and actin (1:5000). Secondary antibodies, anti-rabbit IgG-HRP (1:5000, #7074, CST) or anti-mouse-HRP (1:10000, #12-349, Merck Millipore) in TBST containing 5% milk for 1 h at RT. Bands were detected with SuperSignal™ West Femto Maximum Sensitivity Substrate (#34095, ThermoFisher Scientific) and analysed using the ChemiDoc™ MP (BIO-RAD). Densitometry was performed using Image Lab v6.0.1 (BIO-RAD).

For immunoblotting, patient and donor fibroblasts were grown on 10 cm dishes and lysed in 70 μl Laemmli (BIO_RAD) as described above. Protein concentration was determined using Pierce™ 660 nm Protein Assay Reagent (#22660, ThermoFisher Scientific) mixed with ionic detergent compatibility reagent (#22663, ThermoFisher Scientific). 15 μg of protein was loaded onto 4-20 % gradient gels and samples were immunoblotted with antibodies against phospho-S395-LRRK2 and phospho-T57-eEF2 as above.

### Statistical analysis

Student’s two-tailed t-test or One-way ANOVA followed by post hoc Bonferroni corrections were used as indicated. For patient data, Student’s T-test was used followed by multiple comparison correction with the Benjamini and Hochberg method. Adjusted p-values are shown. The receiver operating characteristic (ROC) curve and Pearson’s correlation analysis were carried out using GraphPad Prism, and p-values calculated by Student’s T-test. For the ROC curve, % sensitivity was calculated from: True positive*100/number of PD individuals and % specificity was calculated from True negatives*100/number of healthy individuals.

### Repository information and ethical permission for human samples

For patient biopsy donors in the Turku University Hospital (TUH) cohort, patient consent was obtained according to the Declaration of Helsinki and approved by an ethical committee at TUH. Patient and control skin fibroblasts from the NINDS cohort were from the NINDS Cell Line Repository (http://ccr.coriell.org/ninds). Patient and control skin fibroblasts from the TNGB cohort were from the “Cell Line and DNA Biobank from Patients Affected by Genetic Diseases”, member of the Telethon Network of Genetic Biobanks (project no. GTB12001), funded by Telethon Italy.

### Data availability

Raw data will be made available upon request.

### Ethical approval

Animal procedures were approved by the Animal Experiment Board in Finland. Patient samples were taken with informed consent and the work was approved by Turku University Hospital (Permission # T175/2014).

## RESULTS

### Rat LRRK2 localizes to and phosphorylates ribosomal fractions

To determine which organelles pathologically active LRRK2-G2019S phosphorylated in brain, we did centrifugal separation of rat brain into fractions and phosphorylated them *in vitro* using purified LRRK2-G2019S. LRRK2-G2019S showed preferential phosphorylation of ribosome-enriched fractions (P3) (Fig. 1A). Moreover, we found that 60 % of endogenous brain LRRK2 co-purified with ribosomes and 20 % with the mitochondrial fraction (P2) (Fig. 1B,C, S1A). We further resolved the ribosomal fraction using sucrose gradient centrifugation and used 40s (eIF2α, eIF2B5 and eIF4G3) and 60s (RPL10a) ribosomal subunit markers to validate the fractionation, as previously (37) (Fig. 1D,E). This established that the bulk of endogenous LRRK2 localizes to the small 40S ribosomal subunits in brain (Fig. 1D).

**Figure 1.**
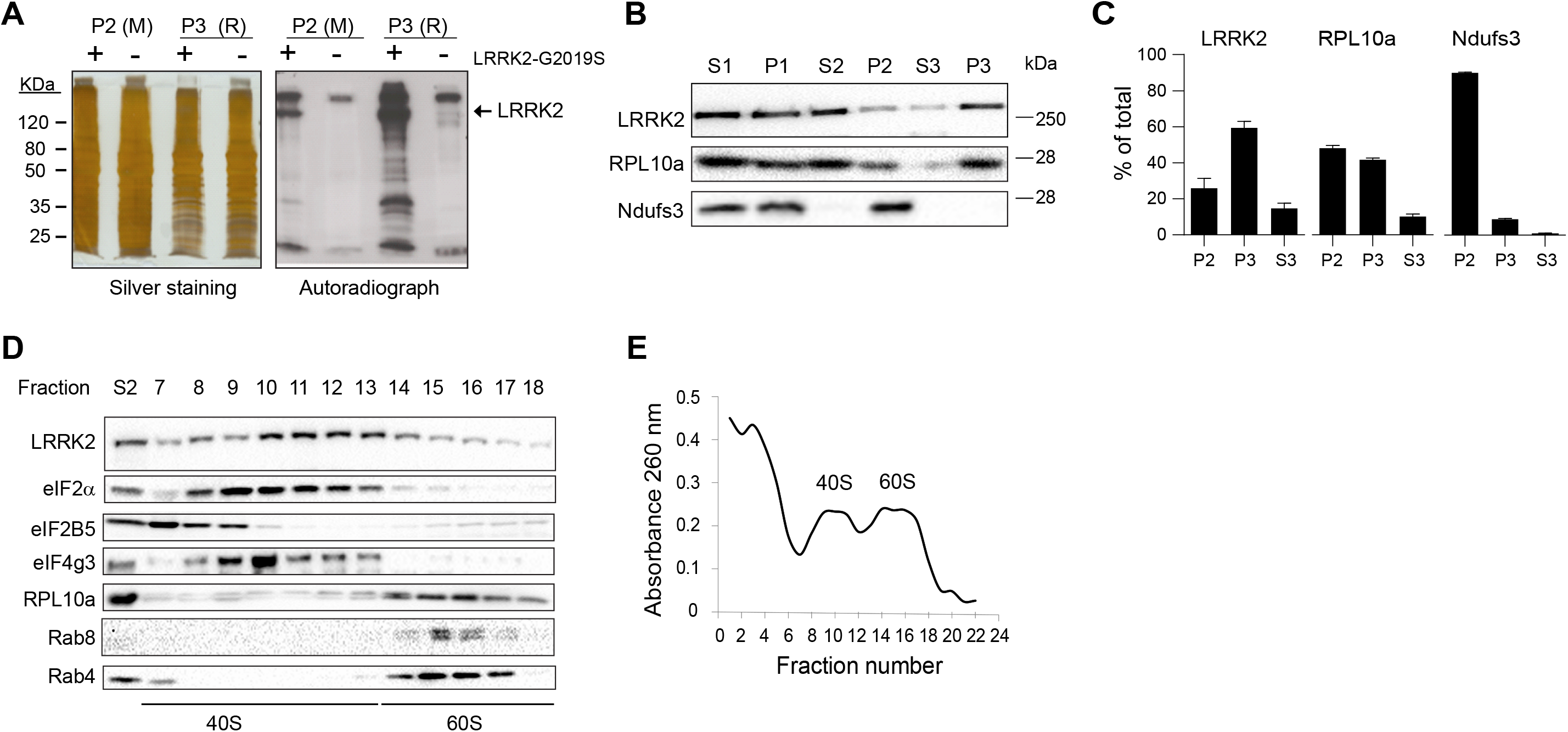
LRRK2 localizes to the 40S ribosomal subunit and phosphorylates ribosomal fractions. (**A)** Mitochondrial (P2 (M)) and ribosomal (P3 (R)) fractions from rat brain were phosphorylated ± LRRK2-G2019S in the presence of [γ-^32^P]-ATP. Silver stained gels and corresponding autoradiographs of phosphorylated fractions are shown. **(B)** Representative immunoblots of rat brain fractions probed with antibodies against LRRK2 and Ndufs3 or RPL10a, to identify mitochondria and ribosomes respectively. **(C)** Quantitative analysis shows relative levels of markers in P2, P3 and S3, expressed as % of total (P2+P3+S3 combined). Mean values ± standard error of the mean S.E.M. from 4 independent repeats are shown. **(D)** Post-mitochondrial supernatant (S2) was separated on 5-50% sucrose gradient and fractions were immunoblotted for ribosomal (eIF2α, eIF2B5, eIF4g3, RPL10a) and endosomal (Rab4, Rab8) markers, as indicated. LRRK2 was enriched at the 40S ribosomal subunit. **(E)** The absorbance at 260 nm for ribosomal fractions shown in D indicates fractions with 40S and 60S ribosomes.

### LRRK2 suppresses RNA translation in neurons

As LRRK2 was enriched at small ribosomal subunits in brain, we tested whether LRRK2 regulated ribosomal function in neurons. Specifically we examined the effect of three structurally independent LRRK2 inhibitors (IN1, GSK-2578215A and MLi-2) (38–40), on *de novo* protein synthesis using non-canonical amino acid labeling (29, 41). We labelled neurons with the Met-analogue L-azido-homo-alanine (AHA), followed by cycloaddition of Alexa-488. Treatment for 1 hour with any one of these three inhibitors increased protein synthesis in cultured hippocampal (and dopaminergic S1B-D) neurons, both in the cell soma and dendrites. Notably, at the dose of inhibitor that reduced both LRRK2 S935 phosphorylation and Rab10-T73 phosphorylation, indicating that LRRK2 was inhibited, protein translation was significantly increased (Fig. 2A-K) (38, 42). As previously reported (43), we found no change in LRRK2 protein levels following 1 hour treatment with LRRK2 inhibitors (S1G-L). As a negative control, we treated cells with anisomycin, an inhibitor of peptidyl transfer. This treatment effectively reduced translation as expected (Fig. 2E; S1E,F).

**Figure 2.**
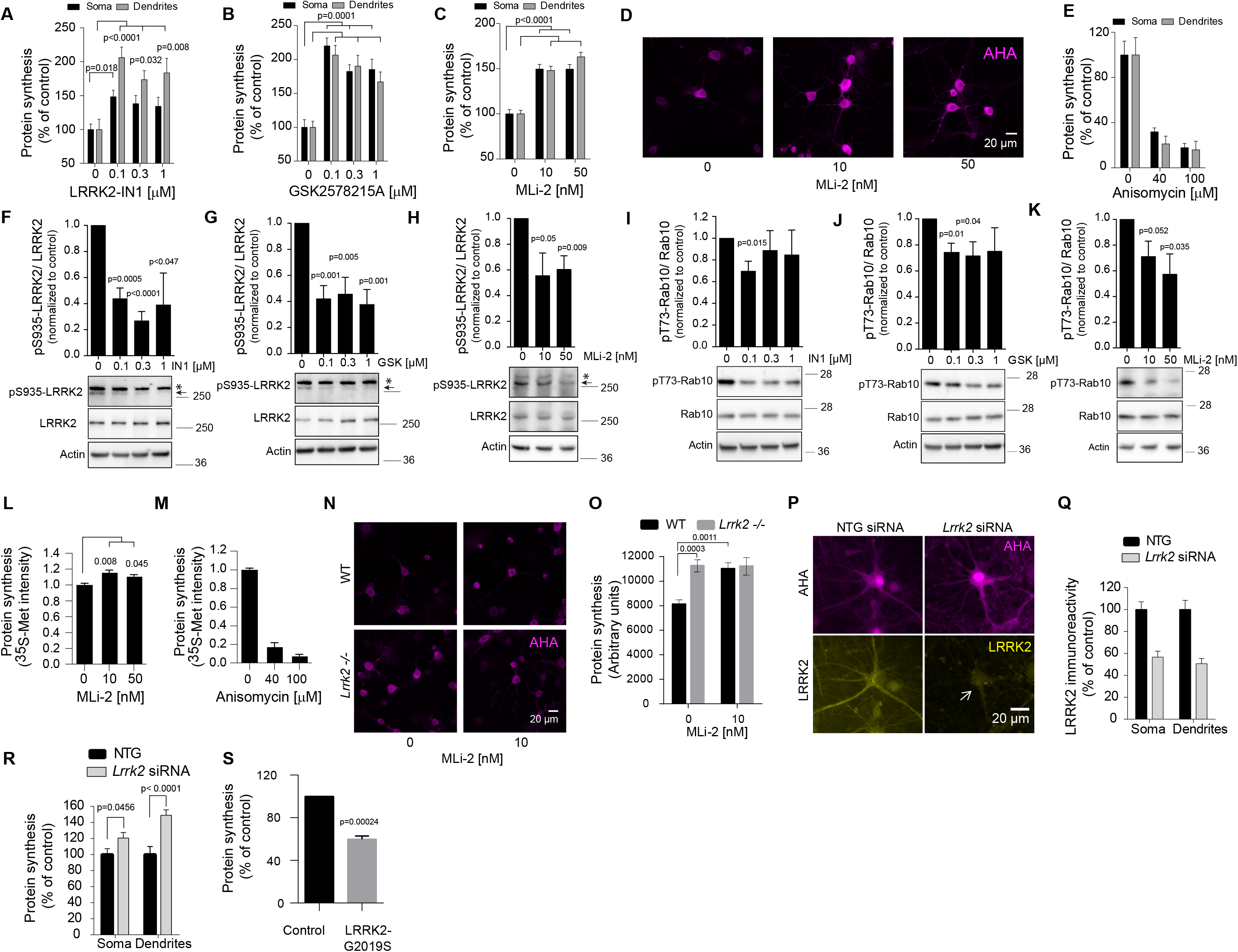
LRRK2 suppresses RNA translation in neurons. **(A-E)** Hippocampal neurons at 20 days in culture were treated for 60 min with LRRK2 inhibitors, anisomycin or DMSO (0.1%), as indicated. **(A-C, E)** Mean fluorescence intensities from soma and dendrites are shown. Mean data from > 20 cells/condition **+/−** S.E.M. are shown. Adjusted p-values determined from one-way ANOVA with Bonferroni post-hoc test are indicated. **(D)** Representative images of AHA-labelling is shown (magenta). **(F-K)** Hippocampal neurons treated as described (A-E), were immunoblotted with antibodies detecting p-S935-LRRK2 and LRRK2 (F-H) or p-T73-Rab10 and Rab10 (I-K), and actin. Means +/− S.E.M. from 4 experiments are shown. p-values were calculated using Student’s t-test. The p-S935-LRRK2 band equivalent to the Rf value of the full length total LRRK2 (indicated by arrowhead) was quantitated. The “*” indicates a non-specific band that is not detected by the total LRRK2 antibody. Molecular weights in kDa are on sides of gels. Quantitative data for LRRK2 and Rab10 are in supplementary material (S1 G-L). **(L, M)** Hippocampal neurons at 20 d were treated with MLi-2 or anisomycin for 90 min, and protein synthesis was measured using ^[35]^S-Met labelling. Quantitative data shows ^[35]^S-Met labelled protein intensity. Mean values +/− S.E.M. relative to control for MLi-2 (n=4) and anisomycin (n=3) are shown. Adjusted p-values determined by one-way ANOVA with Bonferroni post-hoc test are shown. **(N, O)** Protein synthesis was measured using AHA-labelling in WT and *Lrrk2* −/− hippocampal neurons at 16 days in culture, +/− 10 nM MLi-2 for 60 min. Representative images are shown. Means +/− SEM from 52 cells/condition are shown. Adjusted p-values were determined using two-way ANOVA with Bonferroni corrections. **(P-Q)** Hippocampal neurons were transfected with *Lrrk2* siRNA or non-targeting (NTG) siRNA as shown. Representative images are in P. **(Q)** Quantitative data for LRRK2 immunoreactivity is shown. **(R)** The effect of *Lrrk2* knockdown on protein synthesis is shown for 47-64 cells/condition. P-values were determined using Student’s t-test. **(S)** The effect of LRRK2-G2019S on protein synthesis in a cell-free translation system is shown (+/− SEM) from 3 experiments is shown. p-values were determined using Student’s t-test.

We next checked whether LRRK2 inhibitors altered translation measured using classical ^[35]^S-methionine labeling. As low as 10 nM MLi-2 increased protein synthesis in neurons by around 20 % (Fig. 2L), and anisomycin inhibited it (Fig. 2M). This was compared to 50 % increase using AHA, as AHA labelling is more sensitive as it measures from single cell types rather than population analysis (29).

### Neurons from *Lrrk2*−/− mice show increased translation as do neurons after *Lrrk2* knockdown

To validate that effect of LRRK2 inhibitors on protein synthesis were indeed due to LRRK2 inhibition, we measured protein synthesis in neurons isolated from wild-type and *Lrrk2−/−* mice. Consistent with the earlier pharmacological approaches, we found that protein synthesis was significantly increased in cultured neurons from *Lrrk2* knockout mice compared to wild-type littermates (Fig. 2N, O). Notably, treatment with MLi-2 had no further effect on protein synthesis in *Lrrk2−/−* neurons (Fig. 2N, O), further validating that this inhibitor effect on protein synthesis was mediated by LRRK2 inhibition rather than an off-target effect. Finally, we tested the effect of *Lrrk2* knockdown on protein synthesis. Primary hippocampal neurons were transfected with *Lrrk2* siRNA and incubated for 14 days after which endogenous LRRK2 expression reduced by 50% (Fig. 2P, Q). Protein synthesis showed a concomitant increase in *Lrrk2* knockdown cells (Fig. 2R).

### LRRK2-G2019S inhibits translation in a cell free assay

To determine if LRRK2-G2019S acts directly on ribosomes to inhibit translation, we used an *in vitro* translation system where purified ribosomal machinery is used to translate a β-galactosidase reporter in a cell free assay. Addition of recombinant LRRK2-G2019S reduced translation by 40 % (Fig. 2S). This suggests that LRRK2 directly phosphorylates components of the translational machinery to inhibit translation. To distinguish whether LRRK2 regulated regular CAP-dependent translation or IRES-regulated translation, which is important under conditions of oxidative stress (44), we used the bi-cistronic pYIC fluorescence reporter, where CAP- and IRES-dependent translation are simultaneously measured by YFP and CFP respectively. LRRK2 inhibitor increased both CAP and IRES-dependent translation in neurons (S2). In summary, we find using four independent measuring approaches that LRRK2 activity represses protein synthesis in neurons.

### Translation is decreased in rotenone and 6-OH dopamine models of Parkinson’s

We next looked for cellular models of Parkinson’s to test whether translation was impaired. We selected rotenone to model sporadic Parkinson’s disease, as in rodents rotenone reproduces Lewy body hallmarks of sporadic Parkinson’s disease (45), and also because rotenone exposure evokes acute onset of Parkinson’s symptoms in humans and is associated with increased risk to develop Parkinson’s disease (46). We found that treatment of mid brain cultures with rotenone increased LRRK2 activity, as assessed by increased phosphorylation of LRRK2-S935 and of the LRRK2 substrate Rab10 on T73 (Fig. 3A, B, S3A). Consistent with this, rotenone treatment reduced translation by 40% in dopaminergic neurons and this was prevented by LRRK2 inhibitor MLi-2 (Fig. 3C). The same was found in 6-OH dopamine-treated dopaminergic neurons. These results indicate that protein synthesis is repressed in a LRRK2-dependent fashion in cellular models of Parkinson’s disease.

**Figure 3.**
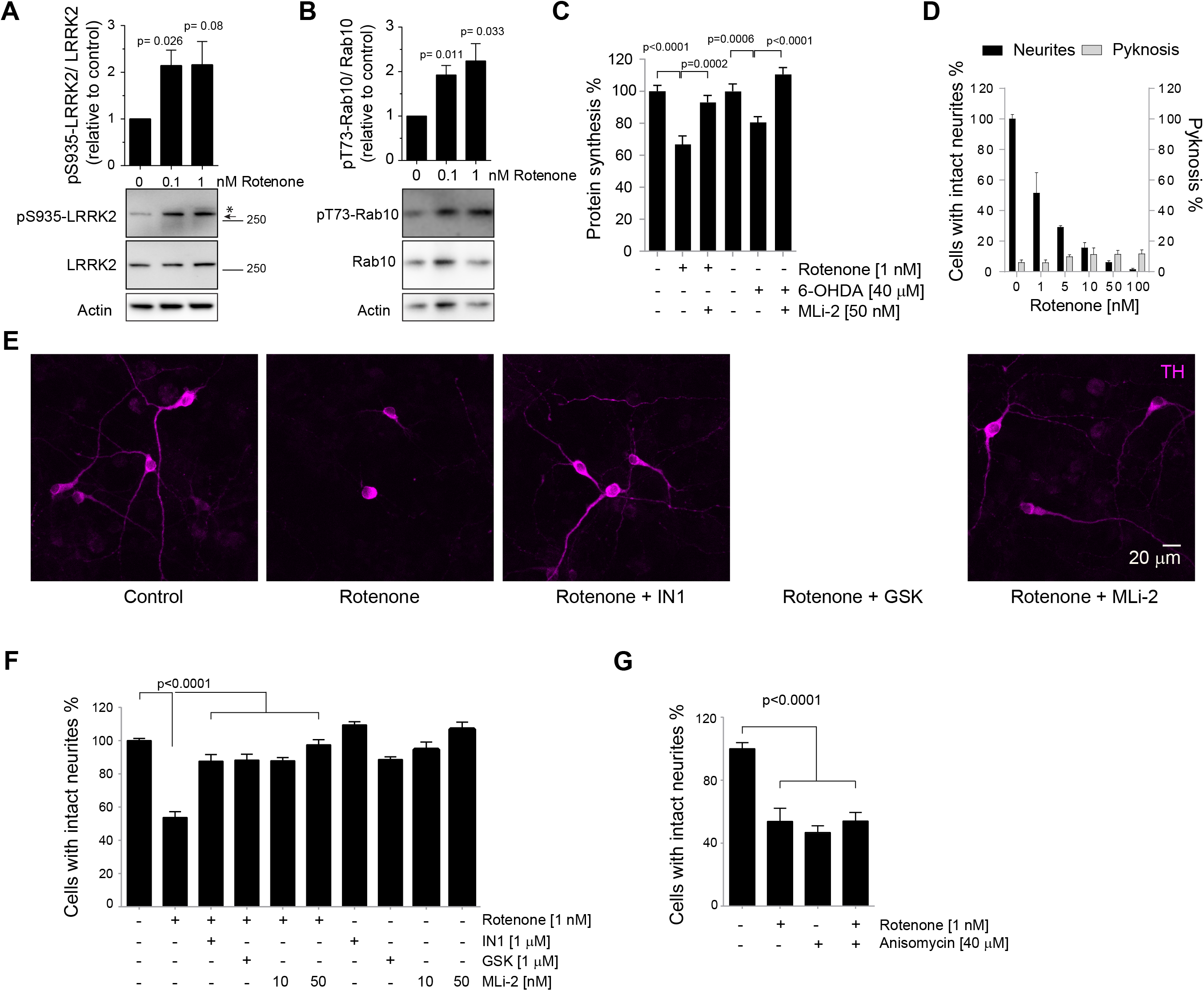
Rotenone activates LRRK2 and induces LRRK2-dependent neuritic atrophy in dopaminergic neurons. (**A, B)** Mid brain cultures were treated with rotenone for 24h. Immunoblotting for p-S935-LRRK2 and LRRK2 (A) or p-T73-Rab10 and Rab10 (B), are shown. Quantitative data +/− S.E.M. is from 3 experiments. p-values were from Student’s t-test. **(C)** TH+ neurons at 3 d were treated with 1 nM rotenone or 40 μM 6-OHDA +/− 50 nM MLi-2. Protein synthesis was measured after 24 h using AHA-labelling. Mean values are from 45-61 cells/condition +/− S.E.M.. p-values from Student’s t-test are indicated. **(D)** The proportion of TH+ neurons with intact neurites (expressed as % of control) or with pyknotic nuclei are shown following 24h rotenone treatment. Mean data +/− standard deviations are shown from two experiments. (**E)** Representative images from F depict TH+ immunostained neurons in magenta. **(F)** Neurite atrophy was measured from TH+ neurons treated with 1 nM rotenone +/− LRRK2-IN1 (1 μM), GSK-2578215A (1 μM) or MLi-2 (50 nM) for 24h. Significance was determined by one-way ANOVA and Bonferroni post-hoc test from 4 experiments. Adjusted p-values are shown. **(G)** TH+ neurons were treated with 40 μM anisomycin or 1 nM rotenone for 24 h as indicated. The proportion of cells with intact neurites is expressed as % of control. Mean values +/− S.E.M. are indicated. Adjusted p-values are from one-way ANOVA and post-hoc Bonfferoni, from 4 experiments.

### Rotenone induces dopaminergic neuron atrophy that is prevented upon LRRK2 inhibition

We next investigated whether LRRK2 activity contributed to neurite wasting or atrophy of dopaminergic neurons, a pathological feature of Parkinson’s disease. Indeed, as low as 1 nM rotenone induced a substantial die back of neurites in dopaminergic neurons in culture without compromising viability (Fig. 3D, E). Neurite atrophy was prevented by LRRK2 inhibitors (IN1, GSK-2578215A or MLi-2; Fig. 3E, F). To determine if atrophy was dependent on repressed translation, we measured the effect of inhibiting translation on neurite integrity. Treatment with either anisomycin or rotenone induced a similar level of neurite atrophy and there was no additional atrophy in neurons treated with anisomycin and rotenone together (Fig. 3G). This shows that actually, repressing protein synthesis is sufficient to disturb neurite integrity and is consistent with our model where rotenone activates LRRK2 leading to repressed translation and compromised integrity of dopaminergic neuron processes.

### Checkpoint regulators of protein synthesis arrest are switched on in the rotenone model

We next tested rotenone *in vivo* in rats. After behavioral testing, brains were extracted and ribosomal fractions were analyzed as earlier (Fig. 1B). Ribosomal and mitochondrial fractions were validated using specific markers; RPL10a for the 60s ribosomal subunit, eIF2α for the 40S subunit and Ndufs3 as a mitochondrial marker (Fig. 4C, D). Interestingly, LRRK2 was enriched in the ribosomal fractions from rotenone treated rats with a concomitant decrease in the remaining supernatant (Fig. 4D). Also notable, expression of the translation repressor 4E-BP1 was increased in brains from rats treated with rotenone (Fig. 4E, F; S3B). These changes indicate that translation is repressed in brain of rotenone-treated rats.

**Figure 4.**
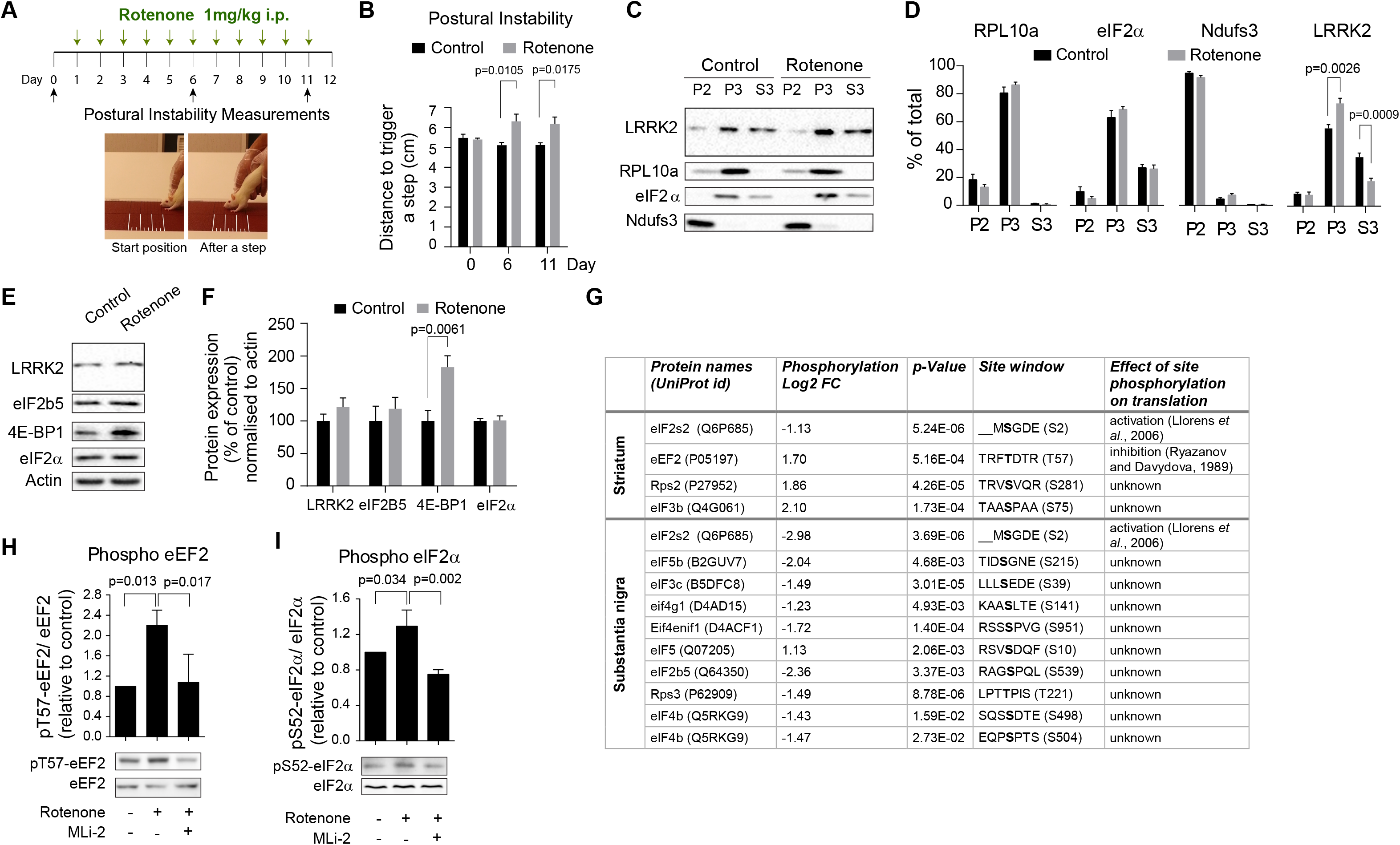
Rotenone regulates phosphorylation of protein synthesis checkpoints. **(A)** Experimental timeline and snapshots of rats undergoing postural instability test. **(B)** Rats were treated with rotenone or vehicle and postural instability was tested from 6 animals per condition. Distance to trigger a step is shown. P-values were determined using Student’s t-test. (**C, D)** Brain fractions from rats after behavioral testing were immunoblotted for ribosomal or mitochondrial markers and for LRRK2. Relative protein levels in P2 (mitochondrial pellet), P3 (ribosomal pellet) and S3 are shown. LRRK2 accumulated at the ribosomal fraction in rotenone-treated rats. Significance was determined using Student’s t-test. **(E)** Immunoblots of brain homogenates from rotenone treated. (**F)** Quantitative data from 6 animals per condition of blots as shown in E. Mean values **+/−** S.E.M. are shown. Significance was determined using Student’s t-test. **(G)** Mass spectrometry analysis of phosphoproteins in substantia nigra and striatum of control (n=12) and rotenone treated (n=14) rats was carried out. Several of the significantly changing phosphoproteins were ribosomal-associated. Phosphorylation fold change (FC) is shown as Log2 scale. p-values (Student’s t test) are indicated. FDR was < 0.05. Aforementioned parameters, as well as protein names, UniProt identifiers and assigned sites are shown. The site window for each phosphosite is indicated with the phosphorylated residue bolded, and effect of phosphorylation on that site is described. **(H, I)** Midbrain cultures were treated with DMSO or 1 nM rotenone −/+ 10 nM MLi-2 for 24 h and lysates were immunoblotted for pT57-eEF2 and eEF2 (H) or pS52-eIF2α and eIF2α (I). Mean data +/− S.E.M. from 3 experiments is shown. Significance was determined by one-way ANOVA and Bonferroni post-hoc test.

We performed a mass spectrometry-based analysis of protein phosphorylation changes in substantia nigra and striatum of control and rotenone-treated rats. This revealed that highly significant phosphorylation changes occurred to regulators of translation initiation and elongation (Fig. 4G). Specifically, in striatum, eIF2s2-S2 (eIF2β), eEF2-T57, eIF3b, and RPS2 phosphorylation was significantly altered. Similarly, eIF2s2-S2, eIF5b, eIF3c, eIF4G1, eIF4E transporter, eIF5, eIF2b5, eIF4b and RPS3 phosphorylation was significantly altered in the substantia nigra. Among these, phosphorylation of eIF2s2 (eIF2β) was decreased on S2 in both regions. eIF2s2 is a part of the trimeric eukaryotic initiation factor 2 (eIF2) complex, that recruits Met-tRNAi as a rate limiting step during translation initiation. Loss of phosphorylation on S2 prevents translation (47). Moreover, phosphorylation of eEF2-T57 was increased. eEF2 translocates nascent peptidyl-tRNAs during elongation. Phosphorylation on T57 inactivates eEF2, thereby reducing elongation (48–51). These phosphorylation changes represent critical checkpoint steps during translation. The phosphorylation changes observed on these sites indicate that protein synthesis is repressed in the striatum and substantia nigra of rotenone treated rats.

We next examined if phosphorylation of translation checkpoint regulators was LRRK2 dependent. Using a phospho-specific antibody against eEF2-T57, we found that rotenone induced phosphorylation of this site was prevented by treatment with LRRK2 inhibitor MLi-2 (Fig. 4H). We also tested phosphorylation of the eIF2 complex on eIF2α–S52, as phosphorylation of this site marks another rate limiting step during translation (52). Rotenone increased eIF2α–S52 phosphorylation, indicating protein synthesis arrest, and this was prevented by treatment with MLi-2 (Fig. 4I, S3C). These data identify that LRRK2 activity induces phosphorylation changes to checkpoint regulators of translation that signal protein synthesis arrest.

### Protein synthesis is reduced in skin cells from sporadic and G2019S Parkinson’s subjects compared to healthy, age-matched individuals

LRRK2 is widely expressed among tissues and by no means confined to brain. We therefore decided to test whether translation was impaired in skin cells from Parkinson’s subjects. We started by measuring translation in skin fibroblasts from Parkinson’s patients and healthy individuals as these have been used to aid mechanistic understanding (53). We obtained from the National Institute of Neurological Disorders (NINDS) repository and from the Telethon Network of Genetics Biobanks (TNGB). Patient demographics, including age, sex, UPDRS and Hoehn & Yahr stage, as well as [^123^I]FP-CIT-SPECT data for striatal dopamine transporter binding are described in Supplementary Table 1. Global protein synthesis was reduced by 40% or greater in cells from sporadic and G2019S patients (Fig. 5A-C). This was reversed upon treatment with LRRK2 inhibitors (Fig. 5B, C), indicating that LRRK2 activity was responsible for reduced protein synthesis not only in G2019S cases, but also in sporadic subjects. To validate these results, we collected additional skin punches from 13 sporadic Parkinson’s patients attending Turku University Hospital (TUH) in Finland and 7, age matched controls. Patients were not fully diagnosed at the time of sampling, but all progressed to have a clinical diagnosis of Parkinson’s disease within 24 months using UK Brain Bank criteria (54). Global protein synthesis was reduced in this TUH patient group, when analyzed alone (p=0.005, size effect = 1.56, power = 0.88; Fig. 5D, black circles) or when combined with NINDS and TNGB cohorts (p<0.0001, size effect = 1.91, power = 0.99; Fig. 5D). These data indicate that repressed protein synthesis provides a specific biomarker readout of Parkinson’s disease from patient cells even at an early stage.

**Figure 5.**
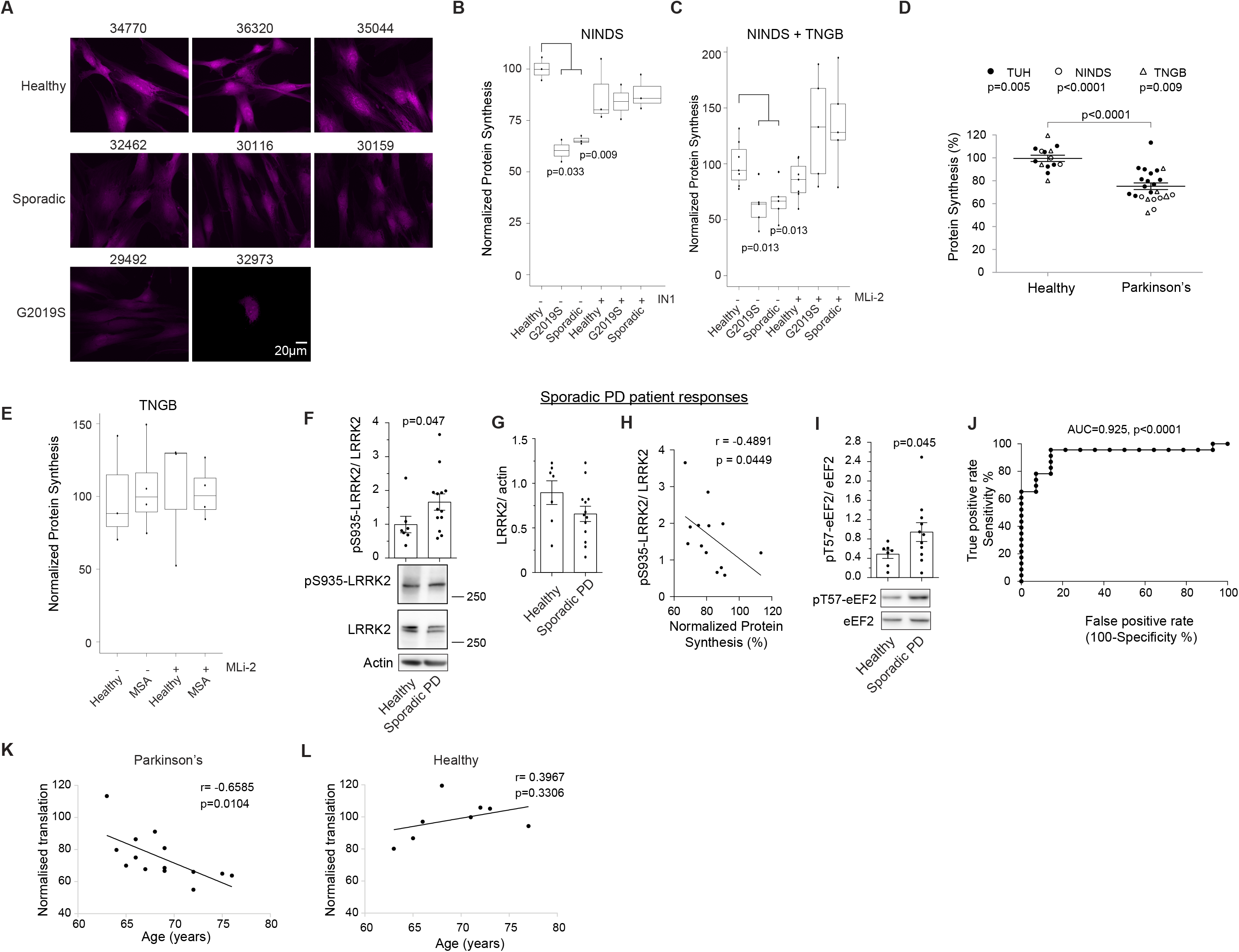
Protein synthesis is reduced in skin fibroblasts from sporadic and G2019S Parkinson’s patients. All patient demographics are in supplementary table 1. **(A)** Representative images of AHA-labeled skin cells from sporadic or G2019S Parkinson’s patients or from healthy controls are shown. NINDS repository numbers are below the panels. (**B)** Quantitative protein synthesis data from healthy, sporadic or G2019S cases +/− LRRK2-IN1 (1μM) treatment are plotted. Adjusted p-values were calculated using T-test followed by Benjamini and Hochberg testing. **(C)** Protein synthesis measured as in B, but with MLi-2 (100 nM) inhibitor. Combined data for samples from NINDs repository, Telethon Network of Genetic Biobanks (TNGB) are shown. **(D)** Cells were isolated from skin biopsies from early stage Parkinson’s patients or healthy volunteers at Turku University Hospital (TUH). Protein synthesis is expressed as mean data +/− S.E.M. Significance was determined using Student’s t-test. **(E)** Protein synthesis data from fibroblasts from Multiple System Atrophy (MSA) patients (n=4) or healthy individuals (n=3). **(F, G)** Cell lysates from sporadic cases were immunoblotted for pS935-LRRK2, total LRRK2 and actin. Mean data shows LRRK2 activity or LRRK2 levels +/− S.E.M.. Significance was determined using Student’s t-test. **(H)** The correlation between protein synthesis and phospho-S935 LRRK2 is shown from patient cells. Pearson’s coefficient (r) and the p-value are indicated on the graph. **(I)** Lysates from F were immunoblotted for pT57-eEF2 and eEF2. Significance was determined using Student’s t-test. **(J)** The receiver operating characteristic (ROC) curve for protein synthesis measured from all patient samples (TUH, NINDS and TNGB) is shown. Area under the curve (AUC) was 92.5%. P-value was determined with Student’s t-test. **(K, L)** Correlation of translation and patient age is shown for patients > 60 yrs. A significant correlation was found between repression of translation and increased age in patients but not in healthy controls.

### Protein synthesis is unchanged in cells from multiple system atrophy patients

Atypical Parkinsonian disorders are mechanistically distinct from Parkinson’s disease but show overlapping symptoms so are difficult to distinguish. Interestingly, we found no evidence that protein synthesis was repressed in MSA patient cells (Fig. 5E). Nor did we find a difference in cells from a patient with progressive supranuclear palsy (data not shown), suggesting that translation deregulation may be specific to Parkinson’s disease.

### LRRK2-S935 phosphorylation correlates with repressed translation in cells from sporadic patients

We next examined whether LRRK2 was activated in skin cells from sporadic patients. LRRK2-S935 phosphorylation increased by 50 % in patient cells compared to healthy donors (Fig. 5F, G). Moreover, there was a correlation between LRRK2-S935 phosphorylation and repressed protein synthesis, consistent with LRRK2 being responsible (Fig. 5H). Also, the elongation checkpoint marker, eEF2-T57 phosphorylation was increased in sporadic patient cells (Fig. 5I), once more indicating repressed translation. Finally, the ROC curve for patient translation data indicated that this measurement provides good predictive power from patient skin cells (Fig. 5J).

### Repression of translation correlates with age in Parkinson’s patients

As age is a major predisposing factor to develop Parkinson’s disease, we measured whether the reduced overall translation response in Parkinson’s patients increased as these patients aged. We found that there was a negative correlation between translation and patient age in individuals older than 60 years (Pearson’s coefficient = −0.6585, p=0.01), but not in younger patients (Fig. 5K, L, S4). Thus, translation became progressively more inhibited in Parkinson’s patients after 60 years of age (Fig. 5K), whereas protein synthesis did not decline in the same way in otherwise healthy, ageing individuals (S4). This fits with the theory that Parkinson’s disease is an accelerated form of ageing, and suggests that reduced translation in Parkinson’s disease may represent accelerated ageing.

In summary, our data shows that LRRK2 is activated in cells from sporadic patients leading to an overall reduction in translation. Measurement of this protein synthesis deficit may provide a means to distinguish between healthy and Parkinson’s individuals.

## DISCUSSION

In this study, we show that LRRK2 represses protein synthesis in animal models of sporadic Parkinson’s disease and in fibroblasts from sporadic and LRRK2-G2019S patients. We use pharmacological inhibition of LRRK2, gene silencing and genetic deletion of *Lrrk2* to demonstrate that repression of protein synthesis is LRRK2-dependent. We identify that known checkpoint regulators of translation are phosphorylated on sites that signal protein synthesis arrest, in animal models and patient cells. Our findings suggest that LRRK2 exerts a repressive regulation of protein synthesis and that this may be a proximal action of LRRK in Parkinson’s disease pathology.

Here we use the rotenone model of Parkinson’s disease as it replicates pathological hallmarks of the sporadic form such as Lewy bodies, postural instability and behavioural deficits that are reversed by L-DOPA (31, 45, 55, 56). Also, exposure to rotenone, a naturally occurring insecticide used in farming, is associated with increased Parkinson’s risk (46), making it a relevant model for sporadic Parkinson’s disease, along with the more commonly used 6-hydroxy dopamine model (57). In both models, we find that LRRK2 is activated and reduces protein synthesis while also inducing atrophy of dopaminergic neurites, an early event in the disease pathology (58). Indeed, we show that merely inhibiting protein synthesis using anisomycin induces dopaminergic neuron atrophy and rotenone has no further effect. This indicates that neurite homeostasis and *de novo* protein synthesis are tightly coupled in dopaminergic neurons and that repression of protein synthesis alone could explain the extent of atrophy obtained with rotenone and 6-hydroxy dopamine (59).

The possibility that LRRK2 may regulate translation has generated interest for some years (17–19, 60). LRRK2 interacts with translational machinery; eIF2C1, EIF2C2 and bifunctional amino-acyl tRNA synthase (16). Also, *EIF2* signalling is severely disrupted in blood from sporadic and G2019S patients alike (61). While these data imply that LRRK2 regulates ribosomal function, direct measurement of translation in mammalian models of Parkinson’s disease or in patient cells, using methods that do not themselves disturb protein homeostasis, have been lacking. Here, we use several methods to measure the effect of LRRK2 on RNA translation. For example, we do two types of metabolic labelling, using either a methionine analogue (AHA), or a methionine isotope (^35^S-Met). Both approaches avoid overexpression. We also measure *in vitro* translation directly in a reconstituted system. Finally, we use a gene reporter to monitor translation in models of sporadic Parkinson’s disease and in patient cells.

We find consistently that translation is repressed by LRRK2 irrespective of the method or disease model used, rotenone and 6-hydroxydopamine models yielding equivalent results.

In contrast to our findings in mammalian cells, human LRRK2-G2019S overexpression in *Drosophila* was shown to increase translation compared to endogenous wild type *Drosophila* LRRK2 (19). This difference may be explained by contextual differences. For example, although human LRRK2-G2019S was expressed in *Drosophila*, it was compared to endogenous *Drosophila* LRRK2 which diverges substantially (16% sequence match) from human LRRK2, and even misses part of the kinase domain (62). Moreover, *Drosophila* ribosomal proteins show poor sequence homology with human, whereas rat ribosomal proteins are 99 % identical to human (63, 64). Lastly, some of the data showing increased translation used reporter overexpression to track protein synthesis (19, 60). Such overexpression overloads cellular resources in particular translational machinery, leading to disturbed homeostasis of cellular machinery and can even trigger promiscuous signalling (65, 66). For this reason, we avoided reliance on overexpression, and rather used amino acid labelling to measure translation. Overall, it is not too surprising that the effect on translation obtained in *Drosophila* models differs from our data in mouse, rat and human cells. It is particularly significant that we find the same effect with endogenous human LRRK2 and human G2019S-LRRK2 in their natural cellular environment, i.e. patient skin cells.

We find that rotenone increases phosphorylation of eIF2α(S52) on a site that inhibits translation at the initiation stage (47, 52). Specifically, phosphorylation of eIF2α(S52) blocks tRNA^Met^ association with the 40S ribosomal subunit, thereby suppressing translation (67–69). Interestingly, increased phosphorylation of eIF2α(S52) is also found in the substantia nigra of Parkinson’s patients(20), and *eiF2α* mRNA is dysregulated (20, 61). We also find that 4E-BP1 is increased by rotenone. This will also inhibit translation by sequestration of eIF4E (70, 71). Finally, we find that eEF2-T57 phosphorylation is increased. This will inhibit its recruitment to ribosomes and stall elongation (48–51). We know from other neurological diseases that disturbed translation causes neurodegeneration, for example in childhood ataxia, where eIF2B5 is mutated (72, 73). In summary, several checkpoints of protein synthesis arrest are switched on in the substantia nigra and striatum of rotenone-treated rats, in rotenone treated dopaminergic neurons and in patient cells. While these phosphorylation changes are not necessarily directly executed by LRRK2, they would appear to be dependent on LRRK2 as they are prevented when LRRK2 is inhibited, suggesting that LRRK2 triggers protein synthesis arrest via these downstream events.

Substantia nigra dopaminergic neurons are selectively vulnerable in Parkinson’s disease. The reason is unknown. One possibility is that they require higher levels of protein synthesis than other cells, making them particularly vulnerable to mechanisms that repress translation, such as described here. Indeed, dopaminergic neurons harbor comparably high numbers of ribosomes (74, 75), believed to reflect the high demand for synthesis of anti-oxidant proteins by these cells (76). Translation rate is directly linked to fidelity of RNA decoding and protein folding (77). Thus by repressing translation, LRRK2 could in theory disturb the cells quality control at many levels, contributing even to α-synuclein oligomer formation, a hallmark of sporadic and LRRK2-G2019S Parkinson’s disease (78). Thus, although LRRK2 expression is widespread among cell types, if pathologically active LRRK2 represses translation, as we posit here, one could envisage that dopaminergic neurons would have a lower threshold of susceptibility, which could in turn impact several aspects of protein homeostasis that are disturbed in Parkinson’s disease.

We demonstrate that LRRK2 is activated and translation is reduced in peripheral cells from patients. While we think this mechanism of LRRK2 is common to all areas where it is expressed, including brain, it raises a question concerning the non-motor symptoms of Parkinson’s disease that occur in the periphery such as olfactory, gastrointestinal, respiratory, skin, sleep, visual and neuropsychiatric dysfunction (2). One could speculate that LRRK2 may contribute for example to disease-associated skin problems e.g. seborrhea and hypo/hyperhidrosis. Moreover, α-synuclein aggregates also appear in patient skin (79), a possible consequence of stalled translation. Moreover, prodromal symptoms of depression and anxiety are already associated with deregulated protein synthesis (80–82), and impaired translation has been factored in sleep disorders (81). Thus, repression of translation by LRRK2 could conceivably contribute to a range of early symptoms.

Of clinical relevance, we find that repression of translation correlates with age in Parkinson’s patients above 60 years, but we detect no decline in translation with ageing in healthy individuals during this time-span. This is consistent with age being the greatest risk factor for Parkinson’s disease and the idea that Parkinson’s is a disease of accelerated ageing (83). Finally, there is a need for a Parkinson’s biomarker (84, 85). Measuring translation from peripheral cells could potentially lead to a biomarker readout. While skin cells are more accessible than cerebral spinal fluid for example, it would also be of interest to examine whether the same mechanism is at play in blood cells. In summary, we identify LRRK2-dependent protein synthesis deficiency in cells from familial and sporadic Parkinson’s disease and in rodent models. Measuring this deficit in peripheral cells from patients shows potential application as a biomarker readout for patient screening and diagnosis.

**Table 1.** Demographic and clinical characteristics of patients and healthy controls -. The source of material either Turku University Hospital (TUH), Telethon network of genetic biobanks (TNGB) or NINDS Coriell repository (NINDS) is indicated. *Unified Parkinson’s Disease rating scale (UPDRS). Gender male (m) and female (f), and age in years when the biopsy was taken, are indicated. Estimated age of onset refers to the age when motor symptoms were first reported. For patients where the relevant information was not available, a hyphen is shown.

| Patient I.D. | Genotype | Source | Gender | Normalized translation | Age | Estimated age of onset | Motor symptom duration (years) | Hoehn & Yahr stage | Motor MDS-UPDRS (Part III)* | DAT binding defect with [123I]FP-CIT |
| --- | --- | --- | --- | --- | --- | --- | --- | --- | --- | --- |
| **Parkinson's patients** |  |  |  |  |  |  |  |  |  |  |
| 1 |  | TUH | m | 76.81 | 60 | 56 | 4 | 2 | 50 | Not performed |
| 2 |  | TUH | m | 80.89 | 69 | 60 | 9 | 2 | 42 | Not performed |
| 3 |  | TUH | m | 81.37 | 59 | 49 | 10 | 3 | 57 | yes |
| 4 |  | TUH | m | 90.04 | 52 | 50 | 1.5 | 1 | 25 | Not performed |
| 5 |  | TUH | m | 79.75 | 64 | 59 | 5 | 2 | 30 | Not performed |
| 6 |  | TUH | m | 69.96 | 65 | 61 | 4 | 2 | 37 | yes |
| 7 |  | TUH | m | 68.54 | 69 | 58 | 11 | 2 | 25 | Not performed |
| 8 |  | TUH | m | 91.15 | 68 | 65 | 3 | 2 | 32 | yes |
| 9 |  | TUH | m | 86.41 | 66 | 64 | 2 | 2 | 32 | yes |
| 10 |  | TUH | m | 89.06 | 49 | 46 | 3 | 1 | 26 | yes |
| 11 |  | TUH | m | 66.88 | 60 | 50 | 10 | 3 | 47 | yes |
| 12 |  | TUH | f | 75.01 | 66 | 56 | 10 | 2 | 15 | yes |
| 13 |  | TUH | m | 113.34 | 63 | 48 | 15 | 3 | 20 | Not performed |
| 29492 | G2019S | NINDS | m | 54.94 | 72 | 58 | 14 | 2 | - | - |
| 32973 | G2019S | NINDS | m | 65.84 | 72 | 62 | 10 | - | - | - |
| 32462 | Sporadic | NINDS | m | 64.94 | 75 | 74 | 1 | 2 | - | - |
| 30116 | Sporadic | NINDS | f | 67.76 | 67 | 43 | 24 | - | - | - |
| 30159 | Sporadic | NINDS | f | 64.08 | 76 | 72 | 4 | 2 | - | - |
| FF0212011 | G2019S | TNGB | m | 52.19 | 47 | 38 | 9 | - | - | - |
| FF0432012 | G2019S | TNGB | m | 90.88 | 57 | 54 | 3 | - | - | - |
| FF0642009 | G2019S | TNGB | f | 64.00 | 58 | 41 | 17 | - | - | - |
| FF0592012 | Sporadic | TNGB | f | 66.69 | 69 | 58 | 11 | - | - | - |
| FF0372011 | Sporadic | TNGB | m | 70.58 | 40 | 38 | 2 | - | - | - |
| **Multiple system atrophy patients** |  |  |  |  |  |  |  |  |  |  |
| FF0582012 |  | TNGB | f | 105.09 | 70 | 65 | 5 | - | - | - |
| FF0662013 |  | TNGB | f | 94.24 | 58 | 56 | 2 | - | - | - |
| FF0522009 |  | TNGB | f | 149.58 | 61 | 50 | 11 | - | - | - |
| FF0212013 |  | TNGB | M | 74.73 | 64 | 61 | 3 | - | - | - |
| **Healthy individuals** |  |  |  |  |  |  |  |  |  |  |
| C1 |  | TUH | f | 92.47 | 57 | - | - | - | - | - |
| C2 |  | TUH | f | 110.49 | 54 | - | - | - | - | - |
| C3 |  | TUH | f | 97.00 | 66 | - | - | - | - | - |
| C4 |  | TUH | m | 108.09 | 58 | - | - | - | - | - |
| C5 |  | TUH | f | 94.04 | 49 | - | - | - | - | - |
| C6 |  | TUH | m | 105.20 | 73 | - | - | - | - | - |
| C7 |  | TUH | f | 86.71 | 65 | - | - | - | - | - |
| 34770 |  | NINDS | m | 105.82 | 72 | - | - | - | - | - |
| 36320 |  | NINDS | f | 99.85 | 71 | - | - | - | - | - |
| 35044 |  | NINDS | m | 94.33 | 77 | - | - | - | - | - |
| F0172014 |  | TNGB | f | 119.53 | 68 | - | - | - | - | - |
| F0492012 |  | TNGB | m | 106.17 | 41 | - | - | - | - | - |
| F0502011 |  | TNGB | f | 80.17 | 63 | - | - | - | - | - |
| F0682013 |  | TNGB | f | 94.12 | 58 | - | - | - | - | - |
| F0342014 |  | TNGB | m | 88.29 | 63 | - | - | - | - | - |
| F0432013 |  | TNGB | m | 70.27 | 62 | - | - | - | - | - |

## Acknowledgements

We thank Susanna Pyökäri, Terhi Hiltula-Maisala, Benjamin Hackl and Lassi Lauren for technical assistance and the Arumäe lab for teaching midbrain cultures. This work was funded by Turku Graduate School of Biomedical Sciences (P.D.), TEKES project # 1817/31/2015 (P.D., P.J., Y.H.), Åbo Akademi University (E.C) and the Michael J. Fox Foundation (P.D.). This project was financed by the Michael J Fox Foundation and TEKES. Salaries were paid by Åbo Akademi University, the University of Turku, the University of Lund and the MATTI graduate school.

## Author roles

EC, PD and DF contributed to conception. EC, PD, DF, YH, PJ, LE contributed to organisation, design and statistical analysis. EC, PD, DF, YH, EG, VS, LS, SP, VK to execution, review and first draft. All authors provided critique.

---

**Fig. S1.**
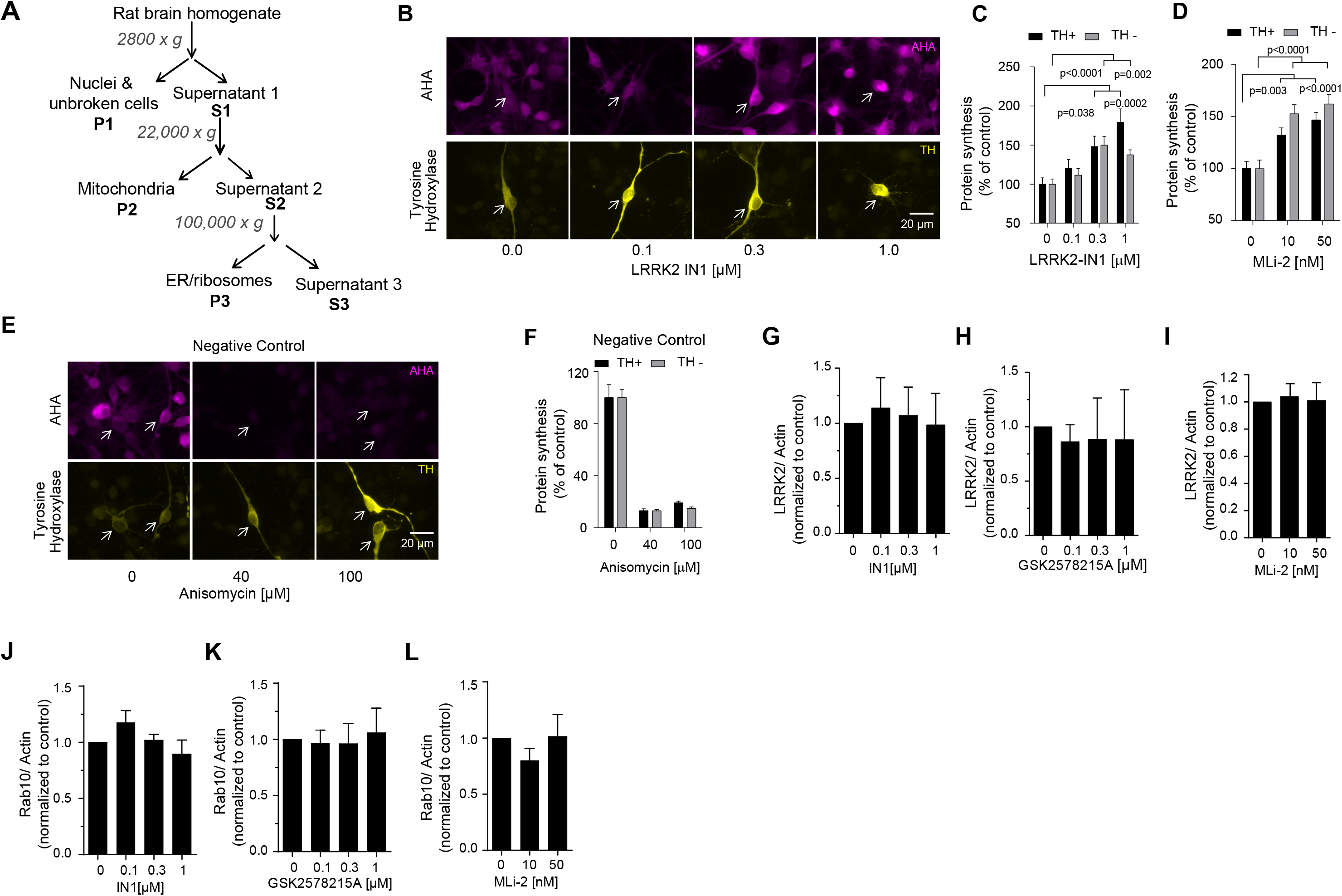
LRRK2 suppresses RNA translation in dopaminergic neurons. **(A)** Schematic depicting mitochondrial and ribosomal isolation steps. **(B-F)** To investigate if LRRK2 regulated protein synthesis in dopaminergic neurons, De novo protein synthesis was measured from midbrain cultures treated with LRRK2-IN1 or MLi-2. Representative images are shown (B, E); AHA-Alexa-488 (magenta) and tyrosine hydroxylase (TH, yellow). Mean soma intensities +/− S.E.M. of AHA-Alexa-488 fluorescence is shown from TH+ and TH− neurons (C,D,F). Data was collected from 10 to 15 neurons per condition for C and F or 35-37 neurons for D. Significance was determined by one-way ANOVA with Bonferroni post-hoc test. **(G-L)** To investigate if LRRK2 kinase inhibitors affected LRRK2 or Rab10 protein levels, we treated hippocampal neurons at DIV20 for 60 minutes with IN1, GSK2578215A and MLi-2 as indicated. LRRK2 levels relative to actin were measured by western blotting. Mean data normalized to DMSO control +/− S.E.M. from 4 independent repeats is shown.

**Fig. S2.**
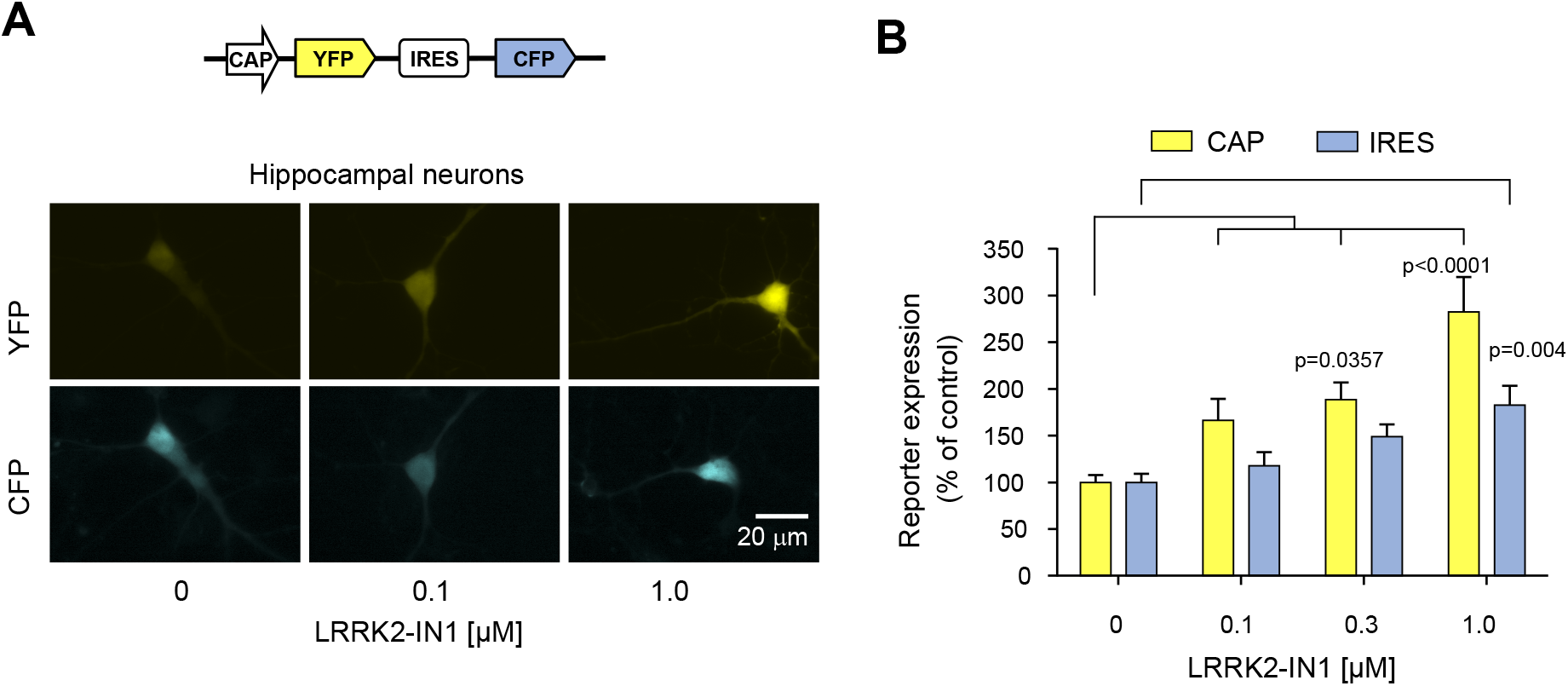
LRRK2 suppresses CAP and IRES-dependent translation in neurons. To study importance of LRRK2 in the regulation of CAP and IRES dependent translation in hippocampal neurons, we expressed the bi-cistronic pYIC reporter in these cells. **(A)** Schematic of the “pYIC” reporter used to measure CAP and IRES dependent translation and representative images of hippocampal neurons transfected at 5 days *in vitro* with it and treated with indicated amounts of LRRK2-IN1 for 48 hours are shown (**B)** Quantitative data showing mean yellow and blue fluorescence output from neurons reporting CAP- and IRES-dependent protein synthesis respectively. Mean data +/− S.E.M. from 16-30 cells per condition are shown. Significance was determined by one-way ANOVA with Bonferroni post-hoc test.

**Fig. S3.**
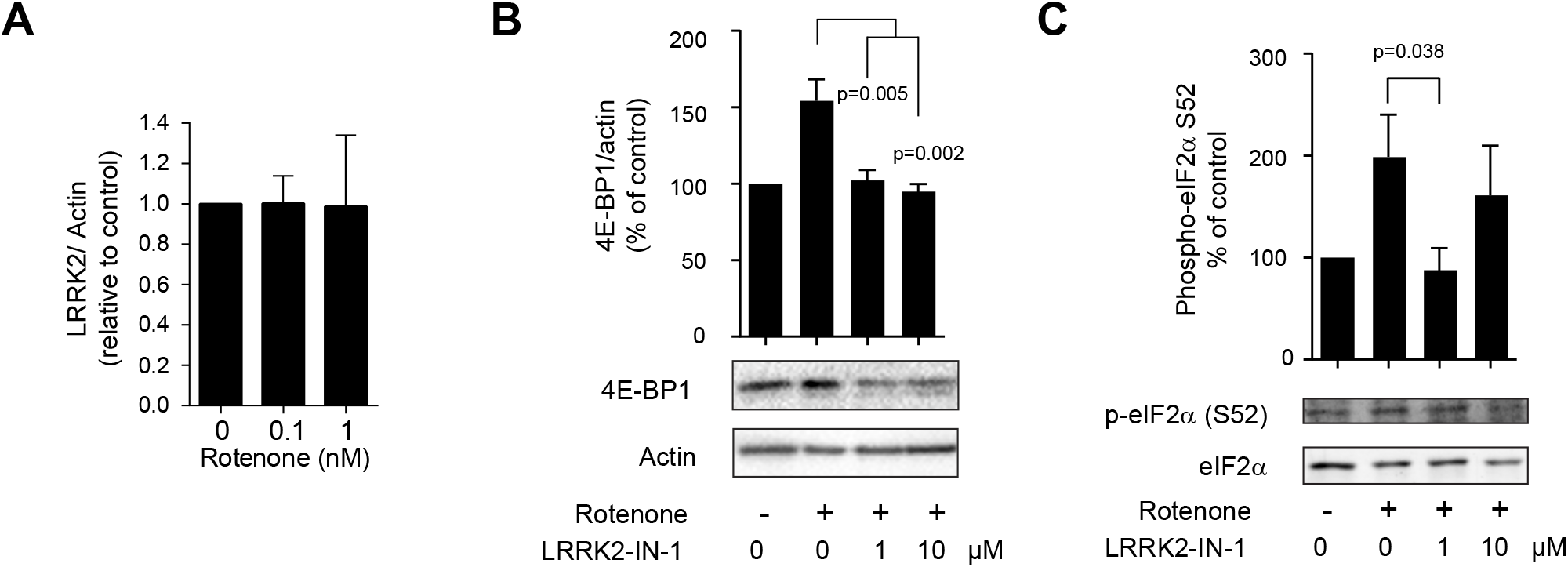
Rotenone treatment induces eIF2α phosphorylation and 4E-BP1 in neurons. **(A)** LRRK2 levels relative to actin in mid brain neurons treated overnight with rotenone. Mean data normalized to DMSO control +/− S.E.M. from 3 independent repeats is shown **(B)** Hippocampal neurons at 18 days post plating were treated with 100 nM rotenone in the presence or absence of LRRK2-IN1. Lysates were immunoblotted for S52-phosphorylated eIF2α (a marker of protein synthesis arrest) and eIF2α. Mean values from 3 independent repeats are shown +/− S.E.M.. P-values were determined using one tailed Student’s t-test. **(C)** Cell lysates from A, were immunoblotted for 4E-BP1. Quantitative data and representative immunoblots are shown. Mean data +/− S.E.M. are shown for 3 experiments. P-values were determined by Student’s t-test.

**Fig. S4.**
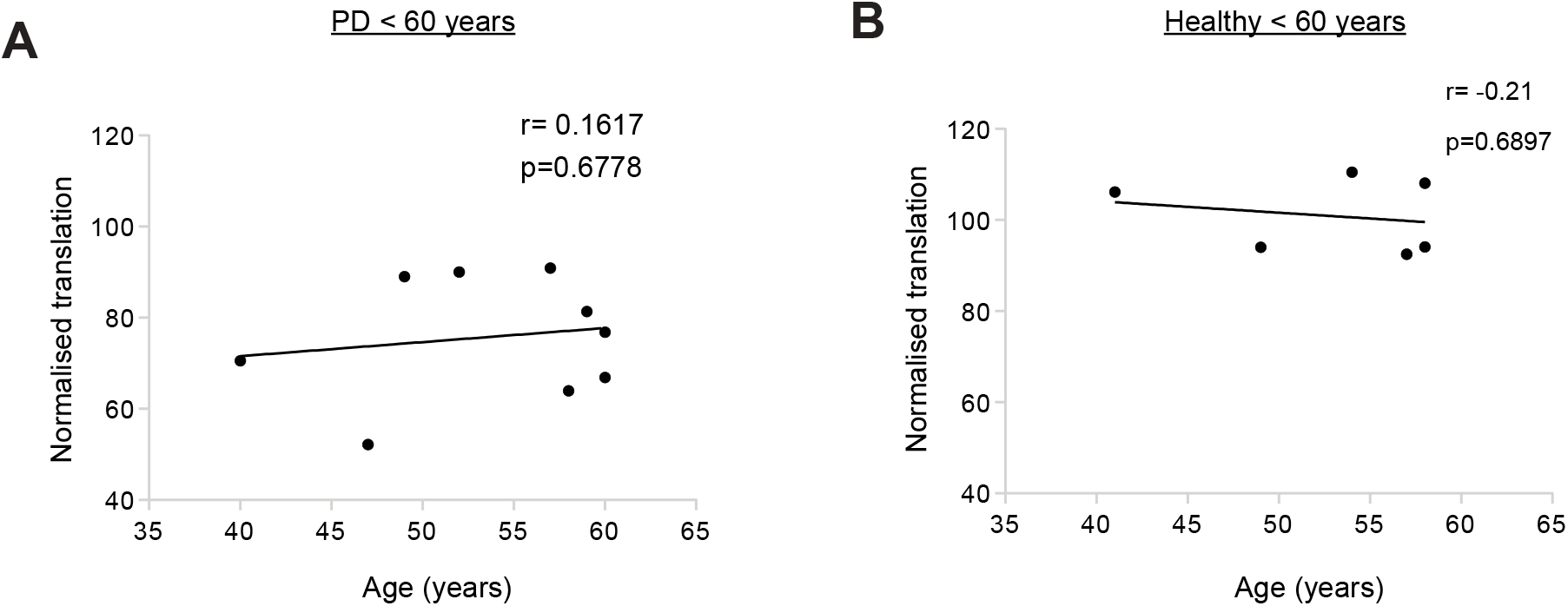
Protein synthesis does not correlate with age at biopsy in fibroblasts obtained from individuals younger than 60 years. **(A,B)** Pearson’s correlation analysis of translation verses age of Parkinson’s patients and healthy individuals younger than 60 years at the time of biopsy. There was no correlation between translation and age in the Parkinson’s disease patient younger than 60 years. Pearson’s coefficients (r) and the p-values (p) are indicated on the graphs.

